# GENETIC DIVERSITY OF *T. B. RHODESIENSE* SERUM RESISTANCE ASSOCIATED GENE IN MALAWIAN ISOLATES

**DOI:** 10.1101/2024.09.07.611819

**Authors:** Peter Nambala, Calorine Claucus, Harry Noyes, Annette MacLeod, Joyce Namulondo, Oscar Nyangiri, Janelisa Musaya, Enock Matovu, Pius Vincent Alibu, Barbara Nerima, Priscilla Chammudzi, Julius Mulindwa, the TrypanoGEN+ Research Group as Members of the H3Africa Consortium

**Affiliations:** Department of Biochemistry and Sports Sciences, College of Natural Sciences, Makerere University, Kampala, Uganda; Kamuzu University of Health Sciences, Malawi-Liverpool-Wellcome Trust Clinical Research Programme, Blantyre, Malawi; Wellcome Centre for Integrative Parasitology, University of Glasgow, Glasgow, United Kingdom; Centre for Genomic Research, University of Liverpool, Liverpool, United Kingdom; Department of Biotechnical and Diagnostic Sciences, College of Veterinary Medicine Animal Resources and Biosecurity, Makerere University, Kampala, Uganda

## Abstract

**Background:** Human African Trypanosomiasis (HAT) is a health burden in most remote areas of Sub-Saharan Africa. Only 2 species of the Trypanosome parasites, namely, *T. b. rhodesiense* and *T. b. gambiense* can establish infection in humans whereas other trypanosome parasites are lysed by human serum APOL-1 protein. The mechanism of *T. b. gambiense* resistance to APOL-1 activity is complex and involves several parasite factors. On the other hand, *T. b. rhodesiense* evades the lytic activity of APOL-1 by intracellular expression of a *Serum Resistance Associated* (SRA) gene that binds to APOL-1 when uptaken by the parasite thereby disabling APOL-1 from causing cellular membrane rupture. APOL-1 has 2 variants, namely, APOL-1 G1 and APOL-1 G2 with the later having mutations on the SRA binding sites which restores APOL-1 lytic activity in parasite lysis assays. This phenomenon remains elusive in clinical setting as limited data is available. In the present study we investigated the genetic diversity of *T. b. rhodesiense* SRA gene and APOL-1 genotypes in Malawian r-HAT clinical phenotypes.

**Methods:** *T. b. rhodesiense* SRA gene from Malawi endemic HAT samples (n= 77) as well as from Zambia and Uganda (n= 13) was amplified by PCR and PCR products were commercially sequenced. APOL-1 variants were identified by restriction fragment length polymorphism (RFLP) after a PCR amplification (n= 61).

**Results and conclusion:** Sequencing data revealed a heterozygosity of the SRA gene within Malawi *T. b. rhodesiense* isolates. Malawian SRA gene was genetically different from some isolates in Uganda and Zambia. Contrary to the current understanding that APOL-1 G2 variants are immune to *T. b. rhodesiense* infection, severe cases of r-HAT in G2 individuals were identified. This study has brought new insight in understanding the determinants of r-HAT severity.

## INTRODUCTION

Human African Trypanosomiasis (HAT) is endemic in Sub-Saharan Africa and causes chronic and acute disease. Most chronic infections are caused by *T. brucei gambiense* whereas, *T. b. rhodesiense* mostly causes an acute disease although some chronic infections have been reported in some disease foci (MacLean et al., 2010). *T. b. rhodesiense* evade the trypanolytic effect of human serum apolipoprotein L1 (APOL-1) by intracellular expression of a *serum resistance associated* (SRA) gene that binds to the N-terminal of APOL-1 and prevents binding to the trypanosome cell wall to cause membrane rupture (Fontaine et al., 2017; Pays et al., 2014; Vanhamme et al., 2003). In some individuals, APOL-1 gene has evolved by having mutations (APOL-1 G1 variants) or deletions (APOL-1 G2 variants) on the binding sites of *T. rhodesiense* SRA. Individuals with APOL-1 G2 variant (Asn388_Tyr389del) may have a five-fold dominance protection against *T. b. rhodesiense* infection (Cooper et al., 2017). Moreover, other experimental results of recombinant APOL1 variants have demonstrated a fivefold protective effect of G2 variants against *T. b. rhodesiense* (Cuypers et al., 2016; Fontaine et al., 2017). APOL-1 G2 variants have also been associated with protection against T. *b. rhodesiense* in Malawi (Kamoto et al., 2019). However, it is not well established whether there are genetic variations of *SRA* gene in endemic *T. b. rhodesiense* in Malawi’s r-HAT foci. Therefore, the aim of this study was to characterize the genetic diversity of *SRA* gene from *T. b. rhodesiense* isolates in Malawi as well as characterize APOL-1 genotypes in individuals from which the *T. b. rhodesiense* were isolated from Rumphi and Nkhotakota r-HAT foci.

## METHODOLOGY

### Samples

We used samples that were collected during active and passive surveillance in Nkhotakota and Rumphi -HAT foci as described (Nambala P et al., 2022). Ethical approval was obtained from Malawi National Health Research Commission to conduct the study. Other samples from Uganda, Zambia and Nkhotakota (collected in 2003) were provided by Annette Macleod’s lab at University of Glasgow where we carried out lab analysis for part of this study.

### Genotyping of *T. b. rhodesiense SRA* Gene

DNA extraction on whole blood in heparin tubes and FTA cards was done using QIAGEN DNeasy Blood and Tissue kit following the manufacturers’ protocol. A PCR of *SRA* gene was done as described (Radwanska et al., 2002), and the PRC products were purified using a ZYMO RESEARCH PCR Purification Kit before proceeding to DNA sequencing. *SRA* gene was sequenced using Sanger sequencing platform to determine genetic variations of the circulating *SRA* gene in Malawi. Sequencing of DNA was outsourced from INQABA Biotechnical Company (South Africa) and Eurofins Genomics (United Kingdom). Furthermore, genotyping of the VSG expression site (VSG-ES) was done as previously described (Radwanska et al., 2002).

### Genotyping of APOL-1 Gene

Determination of *APOL-1* genotype of each *T. b. rhodesiense* infected individual at G1 and G2 haplotype was done using PCR–restriction fragment length polymorphism (RFLP) analysis as previously described (Cooper et al., 2017). This enabled identification of individuals having either APOL-1 G1 or APOL-1 G2 variant *gene*s. APOL-1 G2 variant *gene* is associated with conferring resistance to *T. b. rhodesiense* infection.

### Data Analysis

Analysis of *SRA* gene sequence data was done using QIAGEN CLC Main Workbench v20.0.2. Consensus sequences of Malawi *SRA* gene were obtained in reference to the publicly available *SRA* sequence (Gene ID AF097331.1).

## RESULTS

### Sanger Sequencing Sample Attributes

A total of 90 human blood samples were sequenced by Sanger Platform. 2/90 samples were isolates from Zambia that were collected in 1961, 11/90 were Ugandan isolates with no record of collection date, 3/90 were collected from Nkhotakota in 2003 and 74/90 were collected from Nkhotakota and Rumphi foci from 2016 through 2020. Of the 74 isolates, 46/74 (62.16 %) were from Rumphi foci and 28/74 (37.83 %) were from Nkhotakota foci (**Table 1**).

**Table 1.**
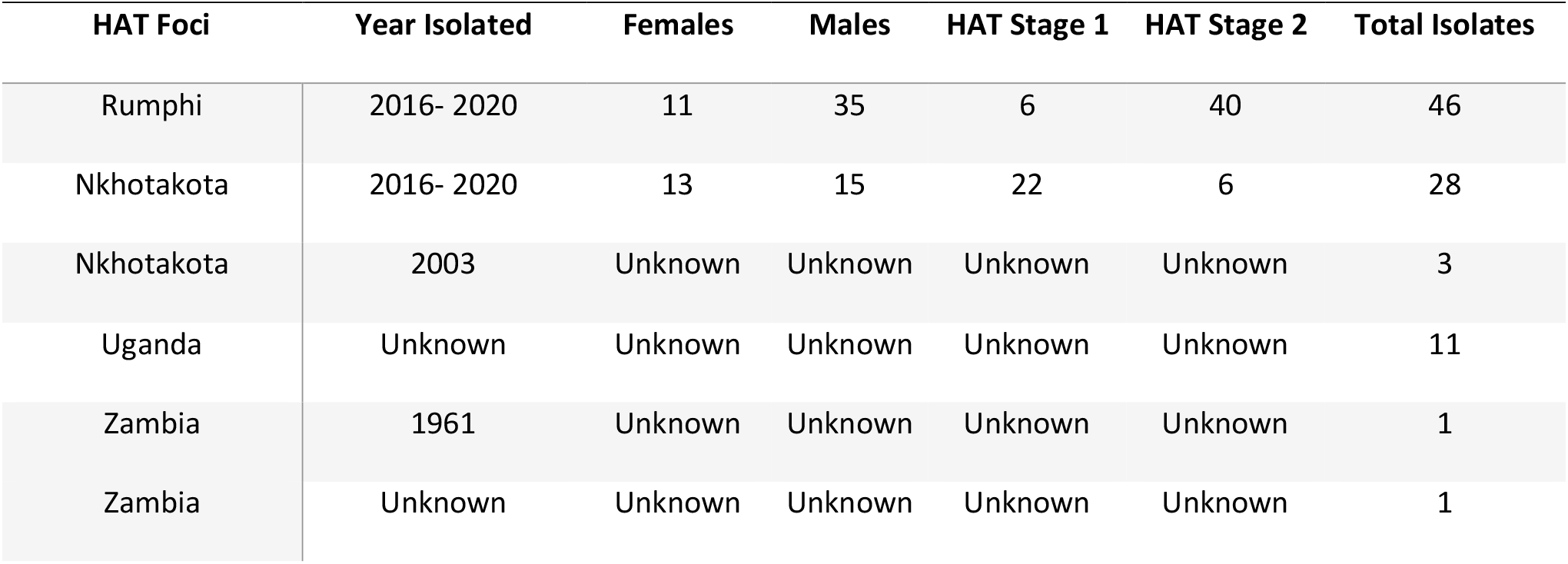
Demographic attributes of individuals from which *T. b. rhodesiense* was isolated.

| HAT Foci | Year Isolated | Females | Males | HAT Stage 1 | HAT Stage 2 | Total Isolates |
| --- | --- | --- | --- | --- | --- | --- |
| Rumphi | 2016- 2020 | 11 | 35 | 6 | 40 | 46 |
| Nkhotakota | 2016- 2020 | 13 | 15 | 22 | 6 | 28 |
| Nkhotakota | 2003 | Unknown | Unknown | Unknown | Unknown | 3 |
| Uganda | Unknown | Unknown | Unknown | Unknown | Unknown | 11 |
| Zambia | 1961 | Unknown | Unknown | Unknown | Unknown | 1 |
| Zambia | Unknown | Unknown | Unknown | Unknown | Unknown | 1 |

### *T. b. rhodesiense* Isolates Circulating in Southern and Eastern Africa have Diverse *SRA*

Gene Since *SRA* gene is an evolutionary mechanism for *T. b. rhodesiense* to establish infection in human (Pays et al., 2014), We next aligned consensus *SRA* gene sequences of isolates from Malawi, Uganda and Zambia and generated a phylogenetic tree. For this, there was genetic distance differences in the *SRA* gene of isolates circulating in Malawi, Zambia and Uganda **(Fig 1)**. Four of Uganda isolates, and one Zambia isolate were phylogenetically the same as the reference *SRA* sequence ID (Z32159.2). The other Zambian isolate was genetically the same as the reference Zambian isolate (AJ345058.1) and different from all isolates from Malawi and Uganda. Eight Ugandan isolates were closely related to some Malawian isolates. Whereas SRA gene of *T. b. rhodesiense* isolates from Nkhotakota that were obtained in 2003 were phylogenetically different from most Malawian SRA genes. This indicate that *T. b rhodesiense* circulating in different HAT foci have genetically distinct *SRA* genes.

### Some Malawian *T. b. rhodesiense* Isolates have Inverted Duplicate SRA Sequences

Generation of consensus sequence revealed that some Malawi isolates had an extended SRA sequence beyond the open reading flame 5’ end of the reference sequence by 364bp (**Fig 2A**). To confirm this extension and rule out Sanger sequencing errors, a qPCR was done on the samples for analysis of high-resolution melting curve as well as SRA gene relative quantity. Differences in nucleotide composition of a gene requires different melting temperature to open the double stranded DNA during denaturation in a PCR process. Therefore, high resolution melting generates sequence-specific melting curves and may reveal variations in the species genotype at the level of a single nucleotide (Chatzidimopoulos et al., 2014).

**Figure 1.**
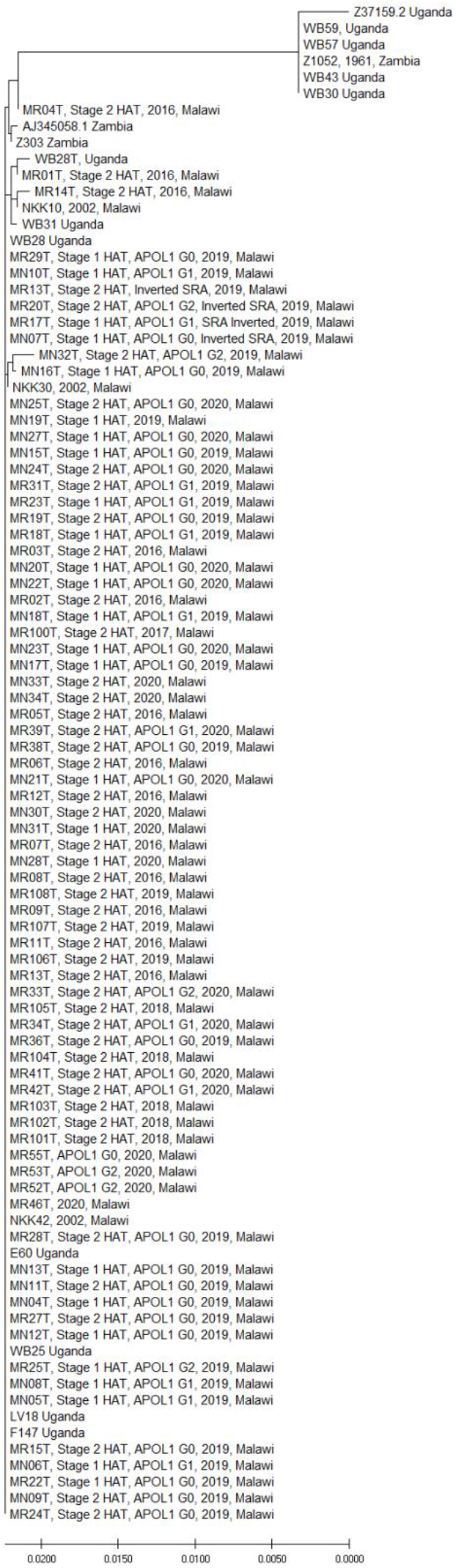
A phylogenetic analysis of SRA gene from T. b. rhodesiense isolates in Malawi, Zambia and Uganda.

**Figure 2.**
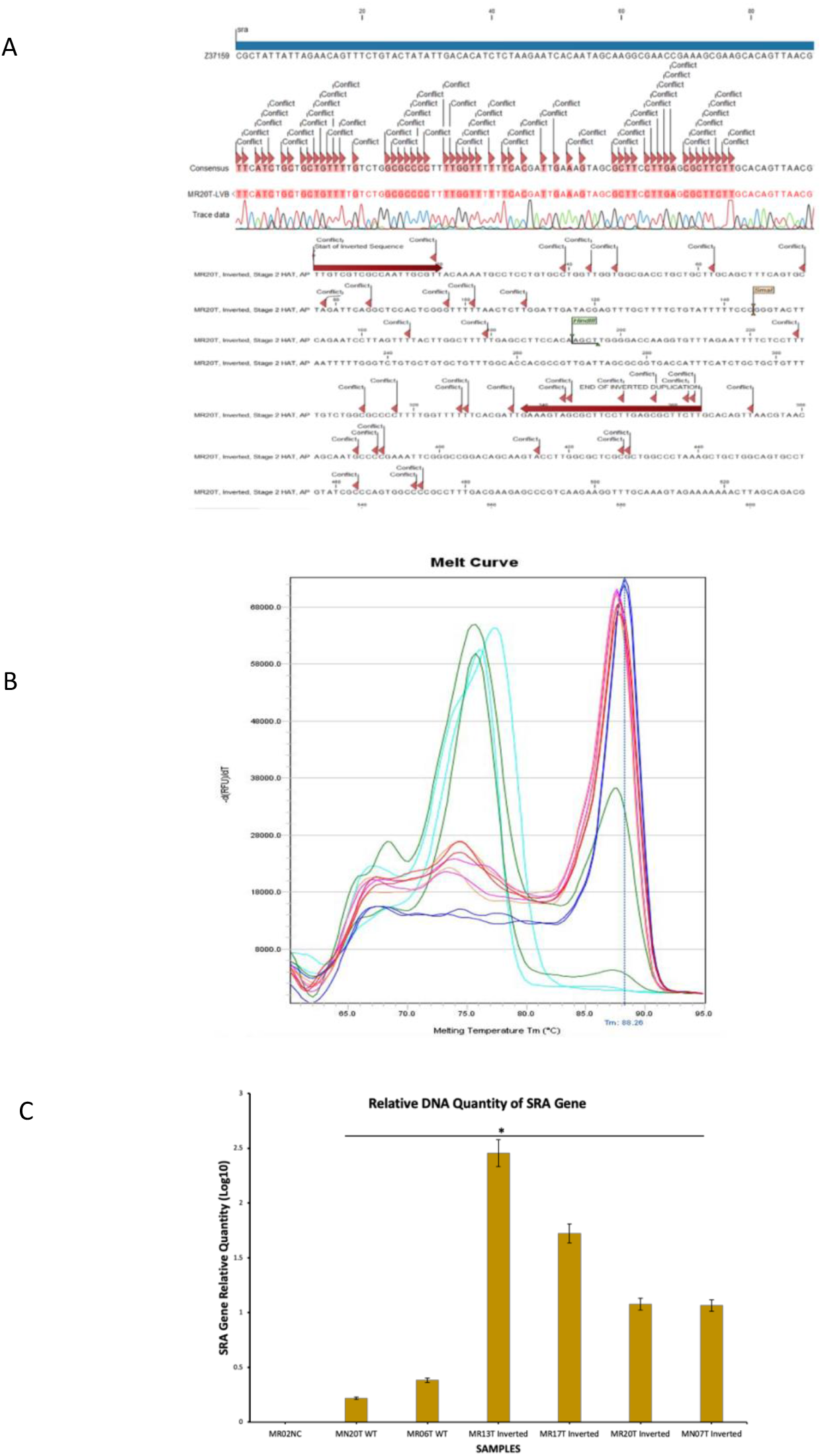
Validation of inverted SRA gene. **A)** Sequence of inverted SRA gene showing the 364bp sequence duplicate extending from the 5’ end of the reference SRA sequence (Z37159.2). **B)** Melting curve of wild type SRA genes (left pick) and inverted duplicated SRA gene (right pick) indicating the differences in melting temperatures between the two variant SRA genes. **C)** A qPCR Comparison of DNA relative quantity between wild type (WT) and inverted SRA genes. *Student test p<0.01

There was a clear difference in the melting curve of wild type SRA genes compared to SRA gene that had an inverted duplicate (**Fig 2B**). Wild type SRA genes had a melting temperature of about 75°C where inverted duplicate SRA genes had a melting temperature of about 88°C. The qPCR relative quantity also showed that there was significantly (p<0.01) more DNA quantity in inverted duplicate SRA gene compared to wild type SRA genes from Malawi (**Fig 2C**). Put together, the results indicate that the nucleotide sequences of wild type SRA genes in Malawi are different from those with inverted sequences.

### The Structure of SRA Protein from Malawi *T. b. rhodesiense* isolates is Different from the Current Reference SRA Protein Structure

The crystal structure of SRA protein has been resolved through Cryo-Electron microscopy (Zoll et al., 2018). The *SRA* protein has three binding sites for APOL-1, the highest binding affinity site is between amino acid I55 and L77 (binding site I)on the SRA N-terminal which interact with APOL-1 C-terminal; the second APOL-1 binding sites are found between Y166 and L172 (binding site II), and the last binding site (binding site III) is between E186 and L201 (Zoll et al., 2018). To determine whether the single nucleotide polymorphisms (SNPs) in Malawi *SRA* gene affected the protein folding, we first converted the SRA DNA sequences from sanger sequencing into amino acid and do amino acid sequence alignment to identify mutations between Malawi, Uganda and Zambia in comparison to the reference sequence (AF097331.1). For this, we identified M75L mutation on the APOL-1 interacting domain in Malawi SRA protein as well as some isolates from Uganda and Zambia (**Fig 3A**). However, there were no mutations on the second and third APOL-1 interacting domains of the SRA protein. High number of amino acid substitutions were found from residue 332 to 358 which is *SRA* protein C-terminal and does not interact with APOL-1 protein (Vanhamme et al., 2003).

**Figure 3.**
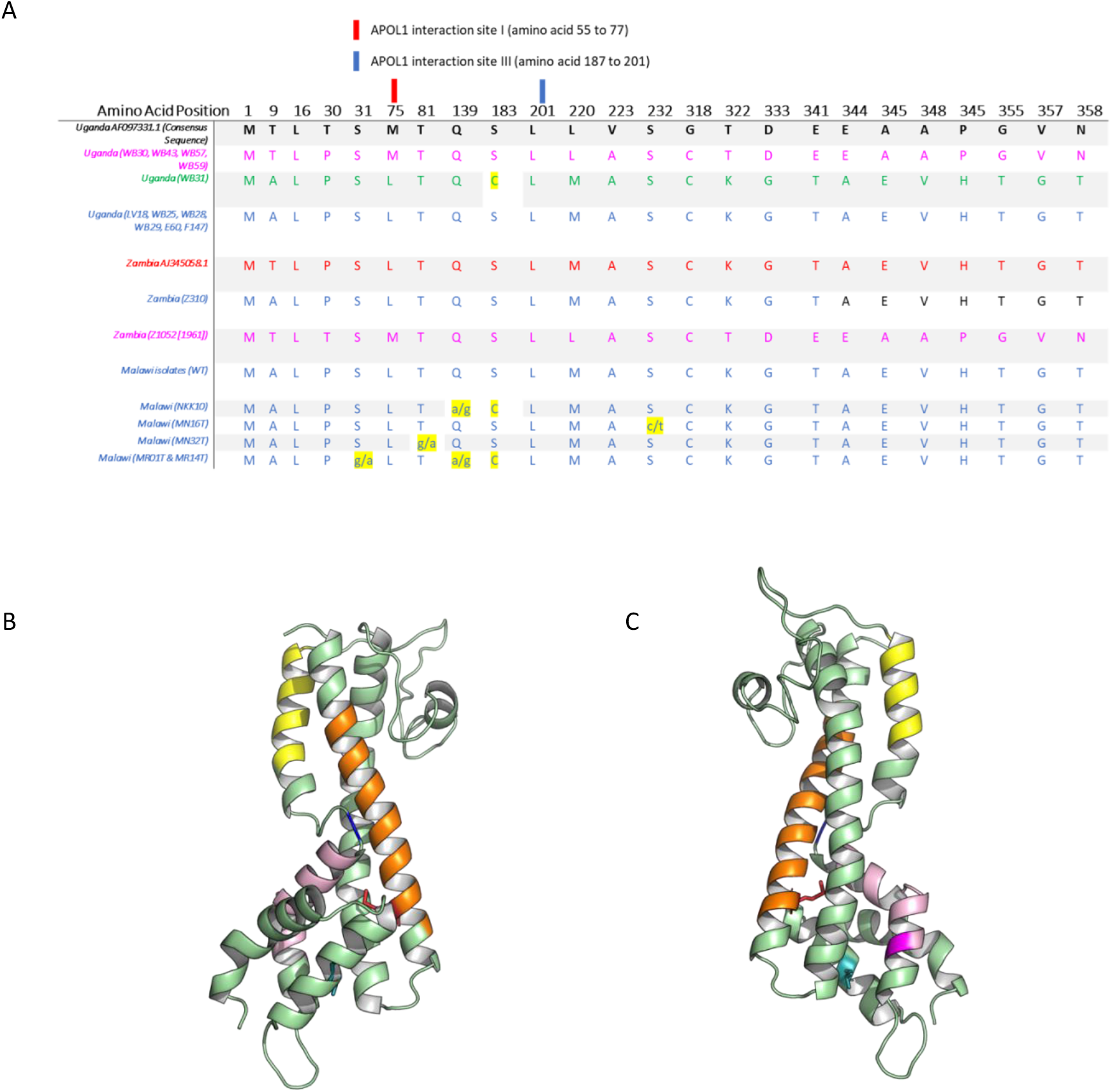
SRA amino acid substitutions and predicted protein model. **A)** Amino acid substitutions on sequenced SRA gene of T. b. rhodesiense isolates from Malawi, Zambia and Uganda in comparison to reference sequence (AF097331.1). Yellow highlights indicate heterozygous mutations or amino acid mutations different from the other isolates. **B)** The structure of Malawi wild type SRA protein model viewed in different angles. Orange colour depicts APOL1 Binding site 1 (55-77) where residue 75 is sticking out; yellow colour is APOL1 Binding site 2 (166-172), and **C)** light pink colour is Binding site 3 (186-201) where residue 201 is the darker pink.

Since we have established that there are mutations on the *SRA* protein domain I, we next checked whether the mutation affect SRA protein structure. The SRA protein sequence were loaded into QIAGEN CLC Main Workbench version 20 in order to identify the best backbone flame for the SRA protein model. The best backbone flame was the one that has been well described (Zoll et al., 2018). The SRA model was further enhanced in BIOVIA Discovery Studio Visualizer. The results shows that the protein folding of Malawi *SRA* changed due to amino acid substitutions in reference to the published *SRA* protein structure (**Fig 3B**). In summary, the *SRA* protein modeling results indicate that variations in SRA genes in *T. b. rhodesiense* isolates from Malawi, Uganda and Zambia result in protein folding changes of the SRA protein.

### APOL-1 G2 Variants were Infected with *T. b. rhodesiense* in Malawi

Since we have established that there is genetic diversity of SRA gene in Malawi’s *T. b. rhodesiense* isolates, next we sought to determine the APOL-1 genotype of individuals from which the isolates were collected. Therefore, 61 whole blood samples from HAT cases were randomly selected and genotyped for APOL1 variants. For this, 22.95% (14/61) of individuals were APOL1 G1 variants and 9.84% (6/61) individuals were APOL1 G2 variants (**Table 2**). Stage 1 and stage 2 HAT was diagnosed in both variants. Interestingly, an individual (MR20T) who had both G1 and G2 variant, was infected with *T. b. rhodesiense* isolate that had an inverted duplicate SRA gene (**Fig 2A**). Our results indicate that APOL1 G2 variant was not protective against *T. b. rhodesiense* isolates in Malawi’s endemic setting contrary to experimental findings in cell cultures.

**Table 2.** Blood samples that were identified as either APOL1 G1 or G2 variants out of the genotyped samples (n=61).

| SAMPLE ID | ApoL1 Variants |  | HAT Stage |
| --- | --- | --- | --- |
|  | G1 Variant | G2 Variant |  |
| MN05T | Yes | No | 1 |
| MN06T | Yes | No | 1 |
| MN08T | Yes | No | 1 |
| MR02T | Yes | No | 2 |
| MR03T | Yes | No | 2 |
| MR09 | Yes | No | 2 |
| MR17T | Yes | No | 1 |
| MR18T | Yes | No | 1 |
| MR23T | Yes | No | 1 |
| MR31T | Yes | No | 2 |
| MR34T | Yes | No | 2 |
| MR39T | Yes | No | 2 |
| MR42T | Yes | No | 2 |
| MR20T | Yes | Yes | 2 |
| MR25T | No | Yes | 1 |
| MR32T | No | Yes | 2 |
| MR33T | No | Yes | 2 |
| MR52T | No | Yes | 2 |
| MR53T | No | Yes | 2 |
| Total Genotyped<br>(n=61) | 14/61 (22.95%) | 6/61 (9.84%) |  |

## DISCUSSION

The *SRA* gene of *T. b. rhodesiense* plays a vital role in rhodesiense HAT as it enables the trypanosome to establish human infections. In this study, we sequenced the *SRA* gene of 28 and 46 endemic isolates from Nkhotakota and Rumphi foci respectively, that were collected from 2016 through 2020. We compared the current endemic isolates of Malawi to those collected in 2003 to identify whether *SRA* gene evolves over time. Our study has shown that, indeed there are genetic differences in circulating *SRA* gene depending on HAT foci. Additionally, we have also found that the SNPs on Malawi *SRA* gene results in changes of *SRA* protein folding in the *APOL1* binding domain. This may have implications on the transmission and pathology of r-HAT as may increases the risk of r-HAT transmission in individuals who were supposed to be resistant to *T. b. rhodesiense* infection.

Furthermore, four *T. b. rhodesiense* (three from Rumphi and one from Nkhotakota) isolates had an inverted duplicate *SRA* gene extending with 364bp from the 5’ end of the wild-type *SRA* sequences. The nucleotide sequence of the 364bp extended part was similar to a section of the *SRA* open reading flame. Melting curve analysis showed that indeed the inverted duplicate *SRA* genes were genetically different from the wild-type *SRA* gene. Future studies should validate the inverted sequence through Cryo-Electron microscopy and how it affects interaction with both wild-type and variant *APOL-1*.

As humans have also evolved in the APOL1 binding domain, it was elusive whether individuals with variant *APOL-1* G2 variants are infected with *T. b. rhodesiense* in endemic settings of Malawi. In this study, we found that 9.84 % of blood samples we genotyped were APOL1 G2 variants and had either stage 1 or stage 2 r-HAT, suggesting that the G2 variant was not protective against *T. b. rhodesiense* infections in endemic r-HAT setting of Malawi. Speculatively, this indicate that the mechanisms of SRA and APOL1 binding are more complex than has been concluded in literature. In this study, we did not perform APOL-1 and SRA protein binding assays using endemic samples, hence this opens an avenue for future studies to determine the effect of protein folding of Malawi SRA protein on APOL-1 binding.

In conclusion, our findings have added to the current understanding on the genetic diversity of *SRA* gene in endemic *T. b. rhodesiense* isolates in Malawi which may impact r-HAT control startegies and clinical pathology. Contrary to the current consensus that APOL-1 G2 variants are protective against *T. b. rhodesiense* infections in cell culture, we found APOL-1 G2 variant individuals that were in either stage 1 or stage 2 of r-HAT indicating that the mechanisms of *SRA* and *APOL-1* protein interaction is more complex than what is currently known. Future studies should explore the mechanisms underpinning this phenomenon.

## Author Contributions

**Peter Nambala:** Conceptualization, Methodology, Investigation, Formal analysis, Writing -original draft. **Harry Noyes:** Conceptualization, Methodology, Formal analysis, Writing -review & editing. **Vincent Pius Alibu:** Conceptualization, Writing -review & editing, Methodology. **Barbara Nerima:** Conceptualization, Writing -review & editing, Methodology. **Joyce Namulondo:** Formal analysis. **Priscilla Chammudzi:** Formal analysis. **Oscar Nyangiri:** Formal analysis. **Enock Matovu:** Conceptualization, Supervision. **Annette MacLeod:** Conceptualization. **Janelisa Musaya:** Conceptualization, Writing -review & editing, Methodology, Supervision, Formal analysis. **Julius Mulindwa:** Conceptualization, Writing -review & editing, Methodology, Formal analysis, Supervision.

## Acknowledgement

We would like to acknowledge Nkhotakota and Rumphi district health offices for their assistance in sample collection.

